# Spike protein disulfide disruption as a potential treatment for SARS-CoV-2

**DOI:** 10.1101/2021.01.02.425099

**Authors:** Andrey M. Grishin, Nataliya V. Dolgova, Shelby Harms, Ingrid J. Pickering, Graham N. George, Darryl Falzarano, Miroslaw Cygler

## Abstract

The coronaviral pandemic is exerting a tremendously detrimental impact on global health, quality of life and the world economy, emphasizing the need for effective medications for current and future coronaviral outbreaks as a complementary approach to vaccines. The Spike protein, responsible for cell receptor binding and viral internalization, possesses multiple disulfide bonds raising the possibility that disulfide-reducing agents might disrupt Spike function, prevent viral entry and serve as effective drugs against SARS-CoV-2. Here we show the first experimental evidence that reagents capable of reducing disulfide bonds can inhibit viral infection in cell-based assays. Molecular dynamics simulations of the Spike receptor-binding domain (RBD) predict increased domain flexibility when the four disulfide bonds of the domain are reduced. This flexibility is particularly prominent for the surface loop, comprised of residues 456-490, which interacts with the Spike cell receptor ACE2. Consistent with this finding, the addition of exogenous disulfide bond reducing agents affects the RBD secondary structure, lowers its melting temperature from 52 to 36-39°C and decreases its binding affinity to ACE2 by two orders of magnitude at 37°C. Finally, the reducing agents dithiothreitol (DTT) and *tris*(2-carboxyethyl)phosphine (TCEP) inhibit viral replication at high µM – low mM levels with a negligible effect on cell viability at these concentrations. The antiviral effect of monothiol-based reductants N-Acetyl-L-cysteine (NAC) and reduced glutathione (GSH) was not observed due to decreases in cell viability. Our research demonstrates the clear potential for medications that disrupt Spike disulfides as broad-spectrum anticoronaviral agents and as a first-line defense against current and future outbreaks.

## Introduction

The world has been significantly affected by the current pandemic of SARS-CoV-2 with close to 80 million total cases and in excess of 1.8 million deceased as of January 2021. Multiple efforts from countries all over the world are being undertaken to find a solution. More than 180 different vaccine candidates are being developed with the anticipation that at least some of them will be efficient (Alturki *et al*., 2020; Krammer, 2020) and the first vaccines have been already administered to risk groups. Clinical trials have been initiated of repurposing current drugs, testing their efficiency against the virus and SARS-induced pneumonia (Oroojalian *et al*., 2020). Novel treatments, ranging from small molecules and biologics (Iacob and Iacob, 2020) to convalescent plasma (Choi, 2020), are being developed.

The family of coronaviridae contains four genera: alpha-, beta-, gamma- and deltacoronaviruses, in which the genera of alpha- and betacoronaviruses includes human pathogens: HCoV-229E, HCoV-NL63, HCoV-HKU1, HCoV-OC43, SARS-CoV, SARS-CoV-2 and MERS-CoV (Zhao *et al*., 2020). The first step in infection of human cells, common to all these viruses, involves the binding of the envelope glycoprotein Spike to a receptor on a cell surface followed by membrane fusion and virus internalization (Yoo *et al*., 1991; Yeager *et al*., 1992). Different coronaviruses use different surface molecules as receptors: SARS-CoV, SARS-CoV-2 and HCoV-NL63 use angiotensin-converting enzyme 2 (ACE2) (Li *et al*., 2003; Hofmann *et al*., 2005; Hoffmann *et al*., 2020); MERS-CoV binds to dipeptidyl peptidase-4 (DPP4) with high affinity (Raj *et al*., 2013) and some sialosides with low affinity (Li *et al*., 2017); HCoV-229E uses aminopeptidase N (hAPN) (Yeager *et al*., 1992); HCoV-OC43 binds 9-O-acetylated sialic acids (Huang *et al*., 2015); while HCoV-HKU1 is internalized through an unknown receptor (Ou *et al*., 2017).

The structures of the Spike protein and its domains have been determined for all of these human pathogens. The functional Spike protein is a homotrimer, in which each protomer is over 1000 amino acids in length (Hogue and Brian, 1986). The Spike protomers consist of two subunits, which in many coronaviruses are cleaved during maturation. The S1 subunit is responsible for receptor binding (Kubo *et al*., 1994) and contains the receptor-binding domain (RBD), while the S2 subunit is responsible for membrane fusion (Yoo *et al*., 1991). The structures of the Spike proteins and their RBDs from different coronaviruses showed that these proteins possessed multiple disulfide (S–S) bonds. For example, the Spike protein of SARS-CoV-2 contains 14 disulfide bonds in well-defined regions (Wrapp *et al*., 2020; Walls *et al*., 2020), that of MERS-CoV contains 11 (Yuan *et al*., 2017) and the Spike of HCoV-229E contains 13 S–S bonds (Li *et al*., 2019). Such an abundance of S–S bridges implies their important structural roles in the formation and stabilization of the proper Spike architecture. This is even more prominent for the relatively small RBD, which has four S–S bonds in SARS-CoV-2 (Lan *et al*., 2020; Shang *et al*., 2020; Wang *et al*., 2020), four in MERS-CoV (Lu *et al*., 2013; Wang *et al*., 2013) and three in HCoV-229E (Wong *et al*., 2017). Mutations of these cysteine residues (Cys) resulted in a significant decrease in expression level or the loss of function. Specifically, individual mutations of seven Cys residues to alanines (Ala) in SARS-CoV RBD resulted in two out of seven mutants losing expression and three other mutants losing the ability to bind ACE2. Only two mutants retained both expression and binding ability of which only in one of these mutants the Cys residue was involved in the formation of a disulfide bond (Wong *et al*., 2004). The knockout of a single S–S bond in the surface loop of the HCoV-229E RBD that interacts with hAPN resulted in the loss of binding (Wong *et al*., 2017). The individual mutations of two conserved Cys residues, forming a disulfide bridge in the S2 domain of SARS-CoV, resulted in a drop of membrane fusion capability to 10% of the wild type and even more for the double mutant (Madu *et al*., 2009). These data suggest that the disulfide-reducing agent could drastically affect the Spike protein structure and function, leading to the inhibition of receptor binding, membrane fusion or both.

Experimental evidence of the usefulness of disulfide-reducing agents in the prevention or mitigation of coronavirus infections is very limited. The proof of principle was demonstrated for SARS-CoV where 3 mM dithiothreitol (DTT) (but not 0.5 mM) could inhibit the viral entry. This result led to a conclusion that SARS-CoV is quite insensitive towards disulfide-reducing agents (Lavillette *et al*., 2006). Recent *in silico* molecular modeling suggested that reduction of only ACE2 disulfide bonds or ACE2 disulfide bonds and SARS-CoV-2 RBD disulfide bonds has a large negative impact on the binding energy, while the reduction of only RBD disulfide bonds does not (Hati and Bhattacharyya, 2020).

Here, we provide the first experimental evidence that the structure of the Spike protein of SARS-CoV-2 is altered by disulfide-reducing agents. As predicted by our molecular dynamics simulations, the structure of the RBD becomes more flexible if not constrained by four S–S bonds. In particular, molecular dynamics shows that the surface loop participating in binding to ACE2 undergoes a very fast conformation opening after the S–S bond between Cys480 and Cys488 is reduced. Experimentally, this flexibility is evidenced by the ability of DTT and *tris*(2-carboxyethyl)phosphine (TCEP) to affect the secondary structure of the RBD and the ability of DTT, TCEP, N-Acetyl-L-cysteine (NAC) and reduced glutathione (GSH) to decrease RBD melting temperature (T_m_). DTT and TCEP could decrease the T_m_ to as low as 36-39°C. As a consequence of increased flexibility and partial melting, the binding constant of RBD to ACE2 of 120 nM increases by 100-200 times in the presence of DTT or TCEP. Finally, the viral propagation in cell-based assays could be inhibited by DTT and TCEP suggesting that disulfide-reducing agents might be useful in the treatment of SARS-CoV-2 infection.

Many recent opinion articles have advocated for NAC as a potential drug against the coronavirus since it is a well-known reducing agent used in medical practice for decades (Rangel-Méndez and Moo-Puc, 2020; Poe and Corn, 2020; De Flora *et al*., 2020). Current clinical trials will provide the ultimate evidence if NAC could be a potential medication for the current and future coronaviruses pandemic.

Our data shows that the SARS-CoV-2 is sensitive to disulfide-reducing agents and compounds with reductive properties could be identified and serve as the first line of defense for the coronaviral outbreak. Broader research to identify best such compounds and proper dosing regimens would have to follow.

## Materials and Methods

### Molecular Dynamics

Molecular dynamics simulations of the Spike RBD domain were conducted in Gromacs 2019 (Pronk *et al*., 2013). For the force field, AMBER99S-ILDN (Lindorff-Larsen *et al*., 2010) was chosen for the Spike RBD with Tip3P water (Jorgensen *et al*., 1983; Neria and Karplus, 1996). The cut-offs for Van der Waals and electrostatic interactions were determined by the Gromacs algorithm. Long-range electrostatics were calculated using the Particle Mesh Ewald approach (Darden *et al*., 1993; Essmann *et al*., 1995). Temperature coupling was done by the v-rescale algorithm (Bussi *et al*., 2007) to maintain the system temperature of 350°K, while the pressure coupling was isotropic and maintained by the Parinello-Rahman algorithm (Parrinello and Rahman, 1981; Nosé, 1984) at 1 atm.

The RBD domain was taken from the ACE2 – Spike RBD protein complex structure PDB ID 6LZG (Wang *et al*., 2020). The RBD structure comprised of residues 333 – 527. The sugar moieties were erased from the PDB files. Three independent simulations were made: 1) RBD with all 4 S–S bonds present as in the X-ray structure (control); 2) RBD with 3 S–S bonds present, while the S–S bond between Cys480 and Cys488 was treated as reduced; 3) RBD with all 4 S–S bonds reduced. The modeling was conducted in a rhombic dodecahedral box in which the proteins were surrounded by a 10 nm thick layer of water. The system consisted of around 44,000 atoms total, 13,700 water molecules and 2 Cl^-^ atoms. The systems were minimized by 1000 steps of the steepest descent method. Minimized systems were subjected to further equilibrium simulations, first for 1 ns with all protein atoms restrained, followed by 10 ns with only backbone atoms restrained. The production simulations were conducted for 2 µs, saving the coordinates every 1 ns.

### Preparation of Compounds

The stock solutions of DL-Dithiothreitol (DTT) (Bioshop, DTT002), *tris*(2-Carboxyethyl)Phosphine (TCEP) (Bioshop; TCE101), Glutathione reduced (GSH) (Bioshop GTH001) and NAC (Sigma-Aldrich; A7250) were titrated to pH 7.6 and filter sterilized through a 0.2 µM filter. Oxidized DTT (*trans*-4,5-Dihydroxy-1,2-dithiane) (Sigma-Aldrich; D3511) was dissolved in DMSO in concentrations up to 2 M or in common buffers (10 mM Phosphate pH 7.5, PBS) in concentrations up to 100 mM.

### Protein Expression and Purification

ACE2 19-615 untagged and Spike RBD 320-541, bearing the C-terminal TEV-His_6_ tag, were cloned into pCEP4 vector, with the N-terminal hemagglutinin signal sequence (KTIIALSYIFCLVFA). The constructs of ACE2 and Spike protein of SARS-CoV-2 were PCR amplified from pCEP4-myc-ACE2, which was a gift from Erik Procko (Addgene plasmid # 141185; http://n2t.net/addgene:141185; RRID:Addgene_141185) and pcDNA3.1-SARS2-Spike -a gift from Fang Li (Addgene plasmid # 145032; http://n2t.net/addgene:145032; RRID:Addgene_145032).

The proteins were expressed in Expi293 cells (Thermo Fisher Scientific) according to the manufacturer’s protocol. The cells, grown in Gibco Expi293 Expression Medium at 37°C at 125 rpm, humidified atmosphere and 8% CO_2_ to a density of 3 million cells/ml, were transfected with 1 µg of plasmid DNA per 1 ml of cell culture and expifectamine® 293 or FectoPro® transfection reagents. Transfection enhancers were added the next day. The media was harvested 4 days after transfection, cleared by centrifugation at 20,000g for 1 hour at +4°C, filtered through a 0.22 µm filter and applied for further purification.

For the purification of ACE2 19-615, the media was applied on HiTrap® Q HP anion exchanger, 5 ml volume (GE Healthcare), preequilibrated with 10 column volumes (CV) of 20 mM Tris-HCl, pH 7.5. After the sample application, the column was washed with 10 CV of 20 mM Tris-HCl, pH 7.5. ACE2 was eluted with 20 mM Tris-HCl, pH 7.5 and NaCl gradient 0 − 0.5 M over 10 CV.

For the purification of RBD, the media was first dialyzed against 20 mM MES, pH 6.1, then applied on HiTrap® SP HP cation exchanger, 5 ml volume (GE Healthcare), preequilibrated with 10 column volumes (CV) of 20 mM MES, pH 6.1. After sample application, the column was washed with 10 CV of 20 mM MES, pH 6.1. RBD was eluted with 20 mM MES, pH 6.1 and NaCl gradient 0-0.5 M over 10 CV.

ACE2 and RBD fractions, eluted from ion exchange, were concentrated and applied on gel-filtration, conducted on Superdex 75 10/300 in 10 mM phosphate buffer, pH 7.5, 50 mM NaCl. The final protein purity was >95% (Figure S1).

### Circular Dichroism (CD)

Proteins were dialyzed overnight against 10 mM phosphate, pH 7.5 and diluted to 0.2 mg/ml. The CD spectra were collected by Chirascan CD spectrophotometer, Applied Photophysics, UK in the range of 195-260 nm in quartz glass cell at room temperature (RT) or 37°C, using 1-nm step size and an acquisition time of 3 s/nm. Data were collected over two accumulations, averaged, smoothed, the background was subtracted. The raw data in millidegrees were converted into molar circular dichroism (Δε).

For spectra acquisition at RT, the proteins were first preincubated with compounds at 2.5 mM final concentration for 1 hr at 37°C, then cooled down to RT. For spectra acquisition at 37°C, the proteins were preincubated with the compounds at 2.5 mM final concentration for 1 hr at 37°C and spectra were acquired at 37°C. RBD with no added compound was also measured at RT without preincubation at 37°C for reference.

For thermal denaturation experiments, the proteins were melted in the temperature range of 20°C - 70°C at a rate of 1°C/min in the presence or absence of the compounds at 2.5 mM final concentration. The CD spectra were acquired every 1°C between 195 and 250 nm with an acquisition time of 1 s/nm. The changes in a CD spectrum as a function of temperature were analyzed with Global3 software (Applied Photophysics) to yield the T_m_ of transitions between protein species observed by CD as the temperature kept increasing. The results represent an average of two independent experiments.

### Thermal Shift Assays

RBD was assayed at a final concentration of 0.5 mg/ml in 10 mM phosphate, 50 mM NaCl, Ph 7.5. Cypro® orange protein fluorescent stain (Thermo Fisher, S6650) was added in a 1:1000 v/v ratio. DTT, DTT oxidized and TCEP were added to yield final concentrations in the range of 0.312 − 5 mM, while NAC and GSH were assayed in the concentration range of 0.625 - 10 mM. Melting was performed in Applied Biosystems StepOne Plus Real-Time PCR amplifier in the temperature range of 20.0-60.0°C in steps of 1°C/min. Fluorescence measurements were taken at every step. The inflection point of a sigmoidal melting curve, calculated in StepOne software, was used as a measure of the melting temperature (T_m_). The results represent an average of two independent experiments.

### Binding Analysis

The affinity constants of ACE2 and RBD were analyzed by microscale thermophoresis, using Monolith instrument (NanoTemper Technologies GmbH). First, RBD was labeled with the RED-NHS 2^nd^ generation Monolith Protein Labeling Kit according to the manufacturer’s instructions (Nanotemper; MO-L011). The fluorescent group was conjugated to RBD through primary amines, while ACE2 was unlabeled. RBD and ACE2 solutions were separately preincubated for 1 hr at 37°C with or without compounds added to the following final concentrations: 2.5 mM DTT, DTT oxidized and TCEP; 10 mM NAC and GSH. Then, RBD was mixed with ACE2 in 20 nM RBD: 1 nM – 50 µM ACE2 ratios in binding buffer 10 mM phosphate, pH 7.5, 500 mM NaCl, 0.1% pluronic acid. The binding experiments were conducted at 37°C according to Monolith’s instruction manual. The results were analyzed in MO.Control and MO.Affinity Analysis software (NanoTemper Technologies GmbH). The results represent an average of at least two independent experiments.

### Coronavirus Infectivity

The ability of SARS-CoV-2 to infect and propagate in eukaryotic cells was performed in the containment level 3 (CL3) facility available at the Vaccine International Disease Organization, International Vaccine Centre (VIDO-InterVac, Saskatoon, SK, Canada). Overall, four disulfide-reducing compounds were tested with 2-fold serial-dilutions at the following final concentrations for DTT and TCEP: 0.15 mM, 0.31 mM, 0.625 mM, 1.25 mM, 2.5 mM, 5 mM, 10 mM and 20 mM; for GSH and NAC: 0.625 mM, 1.25 mM, 2.5 mM, 5 mM, 10 mM, 20 mM, 40 mM, 80 mM and 160 mM.

Vero’76 cells were purchased from the American Type Culture Collection (ATCC, #CRL-1587) and grown to a confluence of 80-90% in Dulbecco’s Modified Eagle Medium (DMEM) (Sigma-Aldrich, D5796, St. Louis, MO, USA) supplemented with 10% fetal bovine serum (FBS) (Thermo Fisher, 16000-044) and 1X Penicillin-Streptomycin (Pen-Strep) (Gibco, 15140148). To evaluate the antiviral potency of these compounds, Vero’76 cells were seeded in DMEM with 10% FBS and 1X Pen-Strep in 96-well flat-bottom tissue culture plates (Millipore Sigma, CLS3595). The SARS-CoV-2/BetaCoV/Canada/ON-VIDO-01/2020 (Sequence available at GISAID: EPI_ISL_413015) virus was diluted in DMEM supplemented with 2% FBS and 1X Pen-Strep to obtain a multiplicity of infection (m.o.i.) of 0.1 (2000 TCID_50_/well). The compounds were serially-diluted in DMEM supplemented with 2% FBS and 1X Pen-Strep, then added individually to both the cells (cells-compound) and the virus (virus-compound) to reach the above-mentioned final concentrations and preincubated for 1 hour at 37°C. After 1 hour, the compounds were removed from the Vero’76 cells (cells-compound) and the pre-incubated virus-compound mixture was added to the cells and incubated for 1 hour at 37°C to allow for viral infection in the presence of the serially-diluted compounds. After 1 hour, the virus-compound mixture was removed and replaced with fresh media containing serially-diluted compounds. The cells were incubated for 48 hours, after which the viral supernatants were harvested and titrated by the TCID_50_ assay, in quadruplicate. The results represent an average of two independent experiments.

In parallel, the serially diluted compounds were added to uninfected cells as a control of cell viability. After 48 hours of incubation cell viability was measured using resazurin (Alamar Blue, Sigma R7017-1G) assay (O’Brien *et al*., 2000). In brief, the growth media was removed and substituted by 100 μl of fresh media mixed with 10 μl of 880 μM resazurin dissolved in PBS. Cells were incubated for 6 hours and the fluorescence was measured at excitation 530 nm and emission 590 nm. The results represent an average of two independent experiments with 3 replicates in each.

To titrate the viral supernatants and determine when viral entry and replication was not blocked by disulfide-reducing compounds, evident by cytopathic effects (CPE), the TCID_50_ assay was performed. To this end, Vero’76 cells were infected with viral supernatants that were serially diluted 10-fold for 1 hour at 37°C. After 1 hour, the virus inoculum was removed and growth medium was added composed of DMEM supplemented with 2% FBS, 1X Pen-Strep and 1 μg/mL L-[(toluene-4-sulphonamido)-2-phenyl] ethyl chloromethyl ketone (TPCK)-trypsin (Sigma-Aldrich). The CPE was observed by visual microscopy at 3 and 5 days post-infection (d.p.i.) for the SARS-CoV-2-induced phenotype, evident by cell death, clustering and rounding of cells. The TCID_50_ titers were determined through the Spearman and Kärber algorithm.

### Statistical Analysis

For the viability assay, the background-corrected fluorescence versus the concentration of compounds was plotted and the 50% cytotoxic concentration (CC_50_) was determined as the concentration of compound that kills 50% of the cells compared to untreated control cells. Viability (%) was plotted, using the untreated cells as the 100% viability reference.

For the viral titration, the TCID_50_ titers were normalized as % of the titer of uninhibited virus and log(%) was plotted against the concentration of compounds. The inhibitory concentration IC_90_ was determined as the concentration of compound that reduces the viral titer by 90% when compared to untreated infected control wells. These results were plotted and IC_90_ and CC_50_ were calculated in GraphPad Prism 8 software, using non-linear four-parameter regression analysis.

## Results

### Disulfide bonds are predicted to rigidify the structure of the RBD domain

To understand whether disulfide bonds are important in the stabilization of the structure of the RBD domain of the Spike protein, we performed 2 µs molecular dynamics simulations of the RBD domain in water using GROMACS software (Pronk *et al*., 2013). The RBD domain (residues 319-541) has four disulfide bonds: C480-C488 is located in the loop interacting with the ACE2 receptor, while three other S–S bonds, C379-C432, C336-C361, C391-C525, are located on the opposite side of the RBD domain and do not participate in ACE2 binding. We performed three independent simulations, first with all four disulfide bonds intact (“RBD”), second with the single disulfide bond between C480-C488 reduced “RBD (−1 SS)” and third with all S–S bonds reduced “RBD (−4 SS)”.

Throughout all three of these simulations, the RBD domain retained its overall fold with the root mean square deviation (RMSD) values plateauing around ∼ 5-6 Å (Figure 1A). The RMSD of RBD increased in a sharp step around 1350 ns. In the meantime, the RMSD for RBD (−1 SS) and RBD (−4 SS) showed a rapid increase in RMSD much earlier in the simulation (Figure 1A). This RMSD increase is related to the conformation transition of the surface loop, spanning residues 456-490, which directly interacts with ACE2. Our simulations show that the disulfide bond C480-C488 stabilizes the loop conformation. For the RBD simulation with the disulfide bridge present, the loop held its conformation for the first ∼1350 ns of the simulation. In RBD (−1 SS) and RBD (−4 SS) simulations, in which this disulfide bond was reduced, the loop started adopting random conformations right from the beginning (200 ns) (Figure 2). This is also reflected in the root mean square fluctuation (RMSF) values for the residues of this loop (Figure 1B), which increased from ∼5 Å to ∼15 Å when the C480-C488 bond was absent.

**Figure 1.**
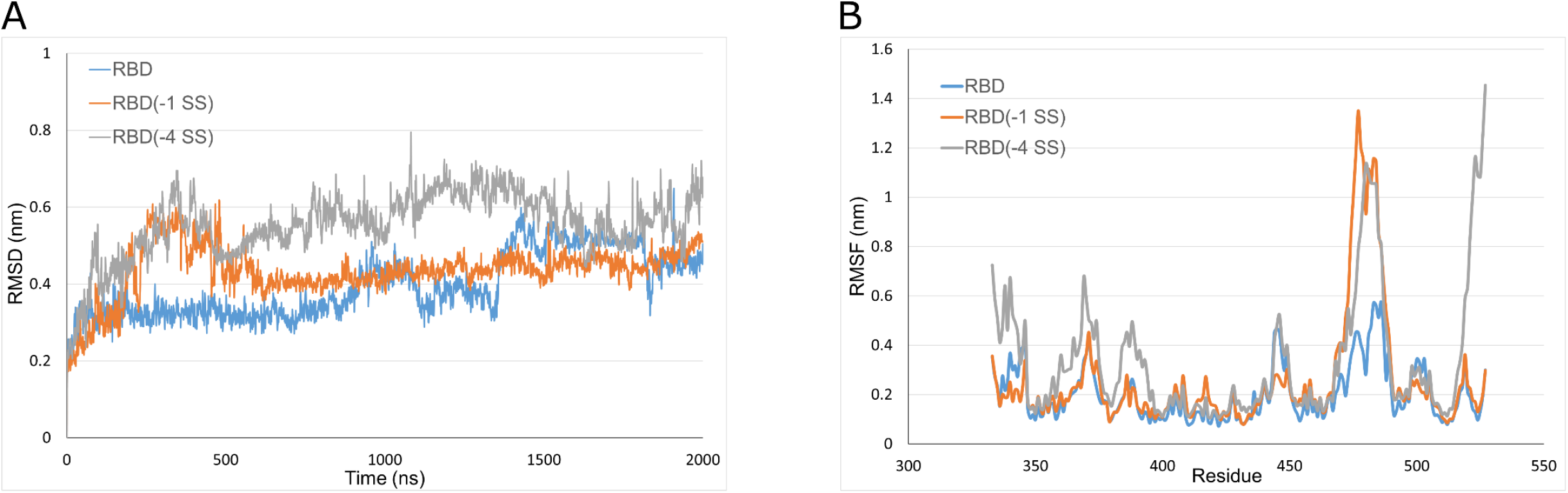
Molecular dynamics of the Spike RBD domain. A) Root mean square deviation (RMSD) of the atomic coordinates (nm) vs. time (ns); B) Root mean square fluctuation (nm) of the residues of the RBD over the first 1 µs of the simulation. The data for RBD with 4 S–S bonds is colored blue; RBD without C480-C488 bond - RBD (−1 SS) – orange; RBD with all 4 bonds reduced - RBD (−4 SS) – grey.

**Figure 2.**
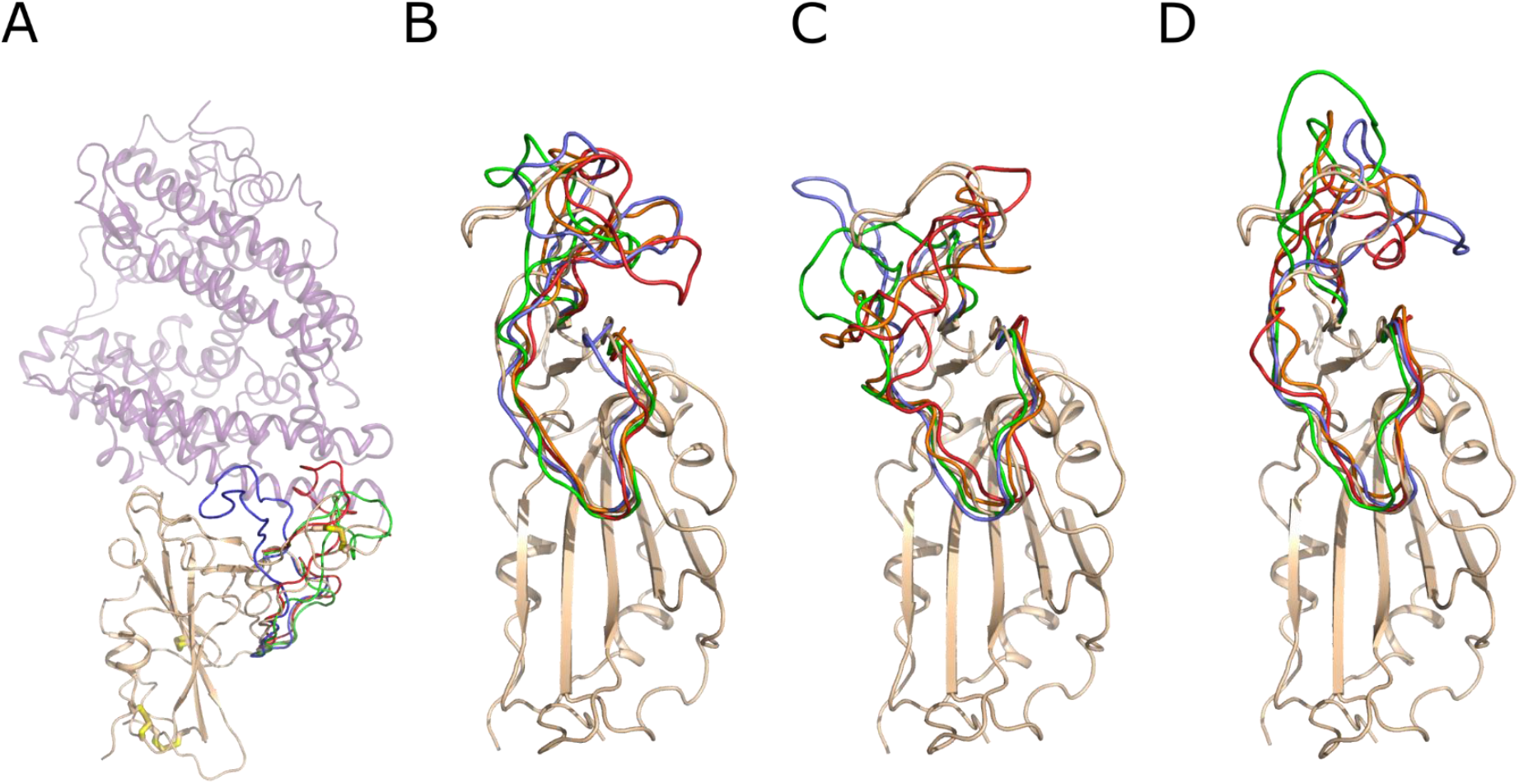
Snapshots of the conformation of the RBD surface loop, residues 456-490, along the molecular dynamics trajectories. A) The crystal structure of the Spike RBD – ACE2 complex. Spike RBD domain is shown as cartoon and colored wheat, while ACE2 is shown as a lilac semi-transparent cartoon. The S–S bonds are shown as wire with white carbon and yellow sulfur atoms. The conformation of the RBD surface loop, residues 456-490, after 250 ns of molecular modeling is shown in green, red and lavender for RBD, RBD (−4 SS) and RBD (−1 SS), respectively. B) Snapshots of conformations of the RBD loop 456-490. The RBD domain is shown as a wheat cartoon. The structure of the loop is shown in lavender, green, orange and red color for 250, 500, 750 and 1000 ns of the modeling trajectory. C) Snapshots of the loop 456-490 conformations of the RBD (−1 SS). D) Snapshots of the loop 456-490 conformations of the RBD (−4 SS). The coloring scheme is identical for B, C and D. The structural deviations of the loop 456-490 are much higher if the Cys480-Cys488 bond is reduced, as shown in C and D.

The other three S–S bonds also contribute to the decreased mobility of protein parts around them, although their absence caused smaller conformational changes than the absence of the C480-C488 disulfide bond. The RMSF graph shows increased mobility of protein parts around residues 340, 370 and 390 when these bonds were absent (Figure 1B). Overall, our simulations predict that S–S bonds are essential for structural stabilization of the RBD architecture.

The surface loop, spanning residues 456-490, undergoes conformational stabilization upon RBD binding to ACE2. The cryo-EM structures of the unbound Spike trimer (Wrapp *et al*., 2020; Walls *et al*., 2020) show that this loop is disordered, while in the crystal structure of RBD complexed to ACE2, this loop is well ordered (Lan *et al*., 2020; Shang *et al*., 2020; Wang *et al*., 2020). The disulfide bond C480-C488, although not positioned on the ACE2 interaction interface, may help in the loop stabilization upon binding to the receptor. Reciprocally, its reduction causes increased loop fluctuations as suggested by molecular dynamics, which may render the adoption of the correct bound conformation more difficult.

### Disulfide-reducing agents destabilize the Spike RBD structure

Since our molecular dynamics results suggested increased flexibility in the RBD domain upon the reduction of disulfide bonds, we investigated whether reagents capable of reducing S–S bonds would introduce structural changes in RBD and ACE2. To this end, we incubated RBD and ACE2 in the presence of DTT, oxidized DTT or TCEP for 1 hour at 37°C and measured their circular dichroism (CD) spectra at 37°C. Oxidized DTT (*trans*-4,5-dihydroxy-1,2-dithiane) served as a control compound, being identical to DTT, except for two thiol groups forming an intramolecular S–S bond. The compounds were added to the final concentration of 2.5 mM, the highest concentration possible without causing a significant increase in the UV light absorption (Micsonai *et al*., 2021).

Incubation of RBD at 37°C did not change its CD spectrum as compared to an RBD sample kept at RT (Figure 3A), suggesting that RBD is stable at human body temperature. However, the addition of DTT and TCEP but not oxidized DTT caused a significant change in the CD spectrum of RBD (Figure 3A), indicating changes in the secondary structure composition, induced by disulfide bond reduction. These changes persisted when the samples were cooled down and analyzed at RT after the preincubation at 37°C (Figure S2). The same experiments conducted for ACE2 (Figure 3B) showed that the addition of DTT or TCEP followed by preincubation and data acquisition at 37°C caused no significant spectral changes, indicating that ACE2 structure is insensitive to exogenous reducing agents. Indeed, ACE2 has only three S–S bonds for a domain of ∼600 residues long and none of these bonds are close to the RBD binding site.

**Figure 3.**
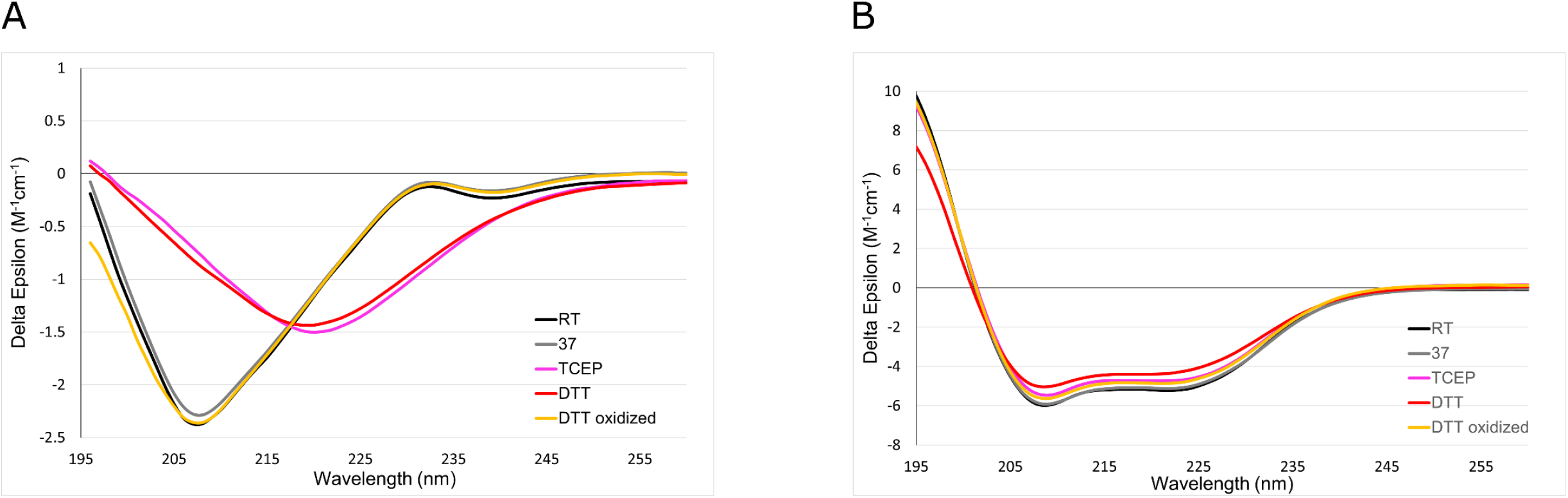
Comparison of circular dichroism spectra of A) RBD and B) ACE2. Both RBD and ACE2 were preincubated for 1 hour at 37°C in the presence of disulfide-reducing agents at 2.5 mM concentration. Spectra were acquired at 37°C immediately after preincubation. The spectrum of RBD and ACE2, never exposed to elevated temperatures and acquired at room temperature (RT), is provided for reference.

Having demonstrated these spectral changes in the presence of DTT or TCEP, we next investigated whether reducing compounds affect the melting temperature (T_m_) of RBD. Indeed, the presence of four S–S bonds in the relatively small RBD domain of 320 residues long should provide a significant contribution to its stability. We conducted a melting experiment of RBD in the presence of 2.5 mM DTT, oxidized DTT and TCEP in which CD spectra were acquired as a function of increasing temperature. RBD exhibited T_m_ of 52.1°C (Table 1), which did not change in the presence of 2.5 mM oxidized DTT (53.3°C). However, a significantly lower T_m_ was measured in the presence of 2.5 mM DTT (39.6°C) and 2.5 mM TCEP (36.3°C). Such low T_m_, observed in the presence of reducing agents, suggests that RBD may partially unfold at normal or elevated body temperatures if the reducing agents are introduced into a human body.

**Table 1.**
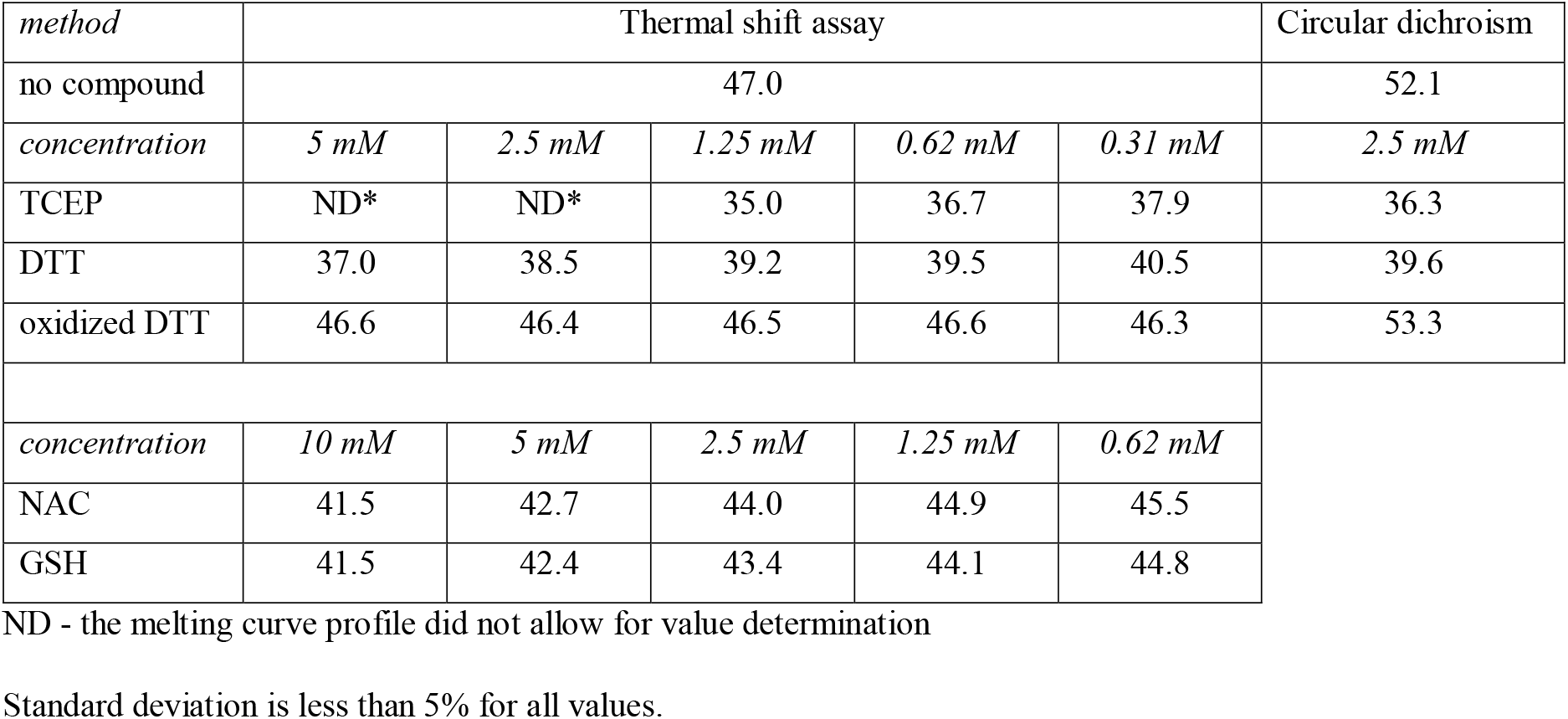
Melting temperatures (T_m_) of RBD in the presence of disulfide-reducing agents at different concentrations Temperatures are expressed in Celsius degrees.

| *method* | Thermal shift assay |  |  |  |  | Circular dichroism |
| --- | --- | --- | --- | --- | --- | --- |
| no compound | 47.0 |  |  |  |  | 52.1 |
| *concentration* | *5 mM* | *2.5 mM* | *1.25 mM* | *0.62 mM* | *0.31 mM* | *2.5 mM* |
| TCEP | ND* | ND* | 35.0 | 36.7 | 37.9 | 36.3 |
| DTT | 37.0 | 38.5 | 39.2 | 39.5 | 40.5 | 39.6 |
| oxidized DTT | 46.6 | 46.4 | 46.5 | 46.6 | 46.3 | 53.3 |
| *concentration* | *10 mM* | *5 mM* | *2.5 mM* | *1.25 mM* | *0.62 mM* |  |
| NAC | 41.5 | 42.7 | 44.0 | 44.9 | 45.5 |  |
| GSH | 41.5 | 42.4 | 43.4 | 44.1 | 44.8 |  |

ND - the melting curve profile did not allow for value determination
Standard deviation is less than 5% for all values.

To broaden our concentration ranges and to explore more disulfide bond reducers, we used a thermal shift assay in which protein unfolding causes a significant increase in the fluorescence of Sypro® orange dye (Senisterra *et al*., 2012). We determined the RBD T_m_ values in the presence of DTT, oxidized DTT and TCEP in the concentration range of 0.31 mM – 5 mM and monothiols NAC and GSH in the range of 0.62 mM - 10 mM (Table 1). High concentrations of TCEP (2.5 mM and 5 mM) precluded accurate T_m_ determination as at these concentrations melting curves lost their typical sigmoidal shape. While all compounds, except oxidized DTT, caused an observable decrease in the T_m_ value of RBD, DTT and TCEP were most potent, while NAC and GSH had a lesser impact than 0.31 mM DTT or TCEP even when added in 10 mM concentrations.

### Disulfide-reducing agents decrease the Spike RBD -ACE2 binding affinity

Since the presence of reducing agents cause the RBD domain to be less stable, we hypothesized that these compounds should negatively affect the binding affinity of RBD to its receptor ACE2. We used microscale thermophoresis (MST) to determine the extent to which 2.5 mM DTT, oxidized DTT or TCEP, as well as 10 mM NAC and GSH, would affect the binding. RBD and ACE2 were preincubated for 1 hr at 37°C in the presence of the reducing agents, after which time the proteins were mixed to yield a constant 20 nM concentration of fluorescently-labeled RBD and a variable concentration of ACE2 in the range of 1 nM – 50 µM. The assays were conducted at 37°C.

The measured affinity of RBD for ACE2 was 120 nM affinity (Table 2, Figure 4), which was not significantly affected by oxidized DTT (170 nM). However, the addition of 2.5 mM TCEP or DTT caused 100-200 times decrease in affinity to 13.3 µM and 21.9 µM, respectively. NAC and GSH also decreased the affinity of the binding between Ace2 and RBD, although the change was not statistically significant even at 10 mM concentration.

**Table 2.**
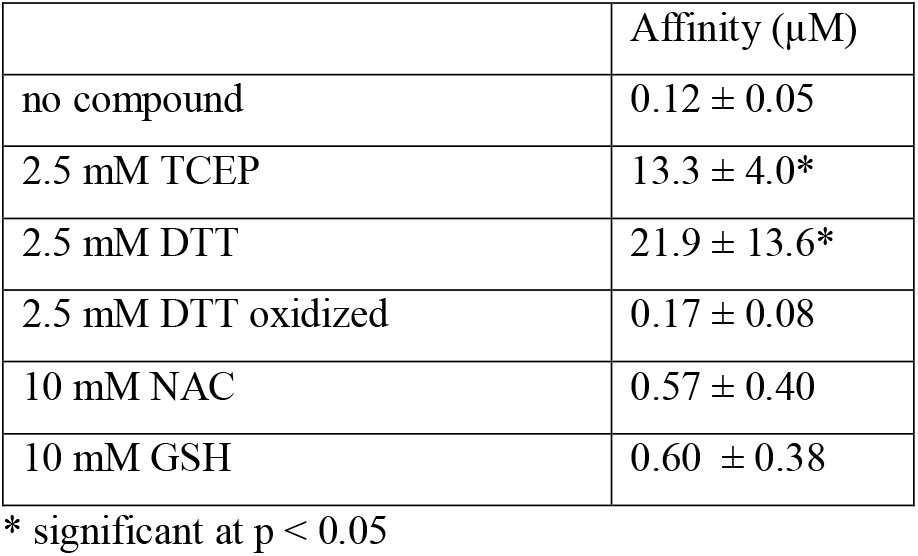
The affinity of the RBD – ACE2 interaction in the presence of disulfide-reducing agents at 37°C determined from microscale thermophoresis (MST).

| | Affinity ($\mu\text{M}$) |
| --- | --- |
| no compound | $0.12 \pm 0.05$ |
| 2.5 mM TCEP | $13.3 \pm 4.0^*$ |
| 2.5 mM DTT | $21.9 \pm 13.6^*$ |
| 2.5 mM DTT oxidized | $0.17 \pm 0.08$ |
| 10 mM NAC | $0.57 \pm 0.40$ |
| 10 mM GSH | $0.60 \pm 0.38$ |

\* significant at $p < 0.05$

**Figure 4.**
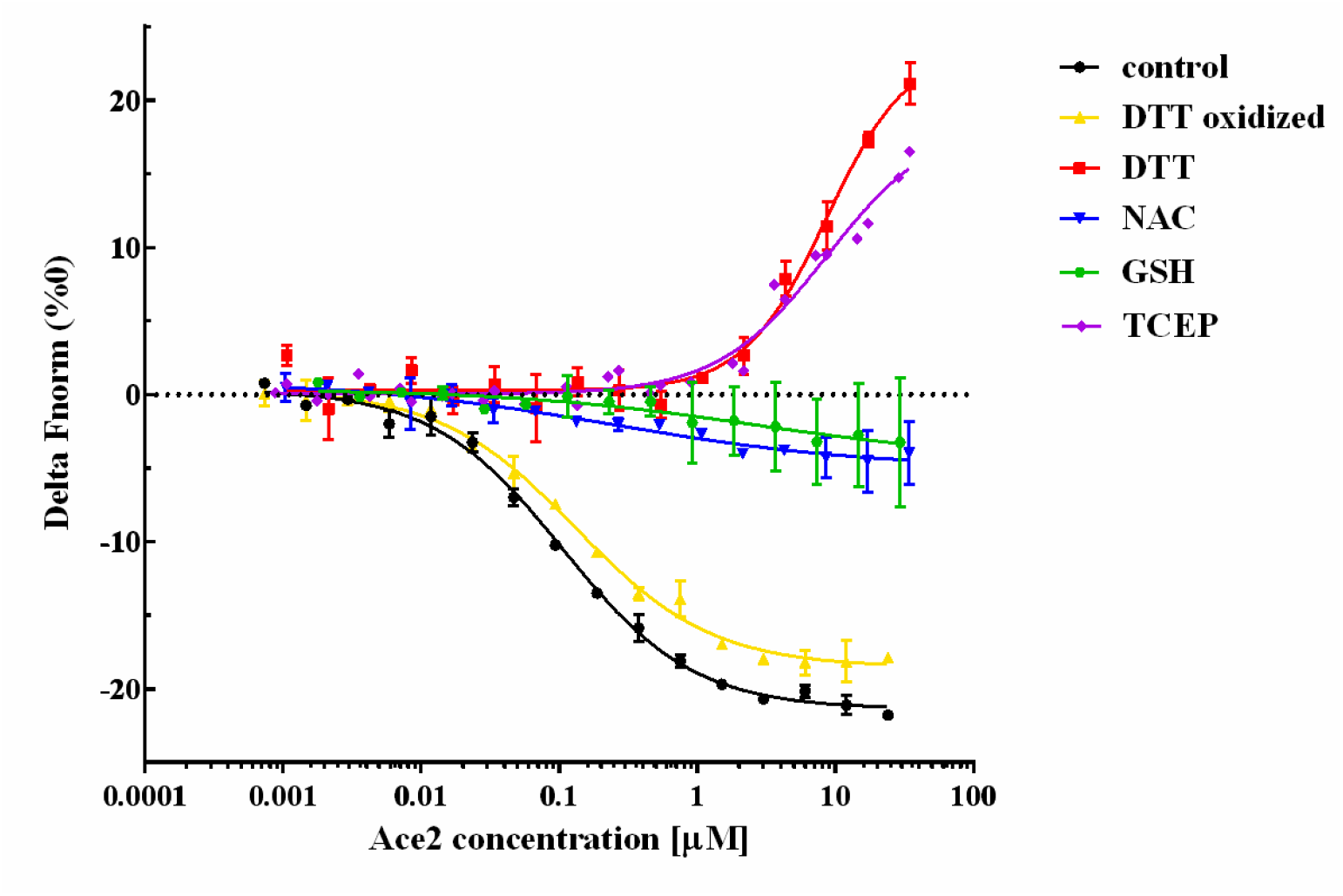
Affinity determination of RBD – ACE2 interaction in the presence of disulfide-reducing compounds at 37°C by microscale thermophoresis (MST). The graph shows the change in baseline-corrected normalized fluorescence (delta Fnorm [‰]) of labeled RBD as a function of ACE2 concentration. The graph shows a sigmoidal binding curve with a plateau at high ACE2 concentrations for oxidized DTT, NAC and GSH. For DTT and TCEP the upper plateau is absent, indicating weaker binding.

### Disulfide-reducing agents prevent coronavirus infectivity in cell-based assays

To evaluate whether reagents capable of reducing S–S bonds inhibit viral replication, we first determined whether the desired concentration range of the respective compounds caused an alteration in cell viability. Cell viability was determined using a resazurin assay (Figure 5), providing a critical concentration 50% (CC_50_) value at which the cell viability is 50% of the control value, and confirmed by phase-contrast microscopy (Figure S3). To avoid interference with the assay caused by redox-active compounds, the cellular medium was replaced after 48 hours of incubation with a fresh medium with resazurin but without reducing agents. DTT is the most toxic of the tested reducing agents (Figure 5) with the critical concentration 50% (CC_50_) ∼2.4 mM. Monothiols NAC and GSH show the lowest toxicity with CC_50_ at ∼50 mM, while TCEP CC_50_ can be fit to ∼7 mM, although ∼30% of cells are viable at TCEP concentrations exceeding 20 mM, and ∼40% of cells survive in 80-100 mM of NAC.

**Figure 5.**
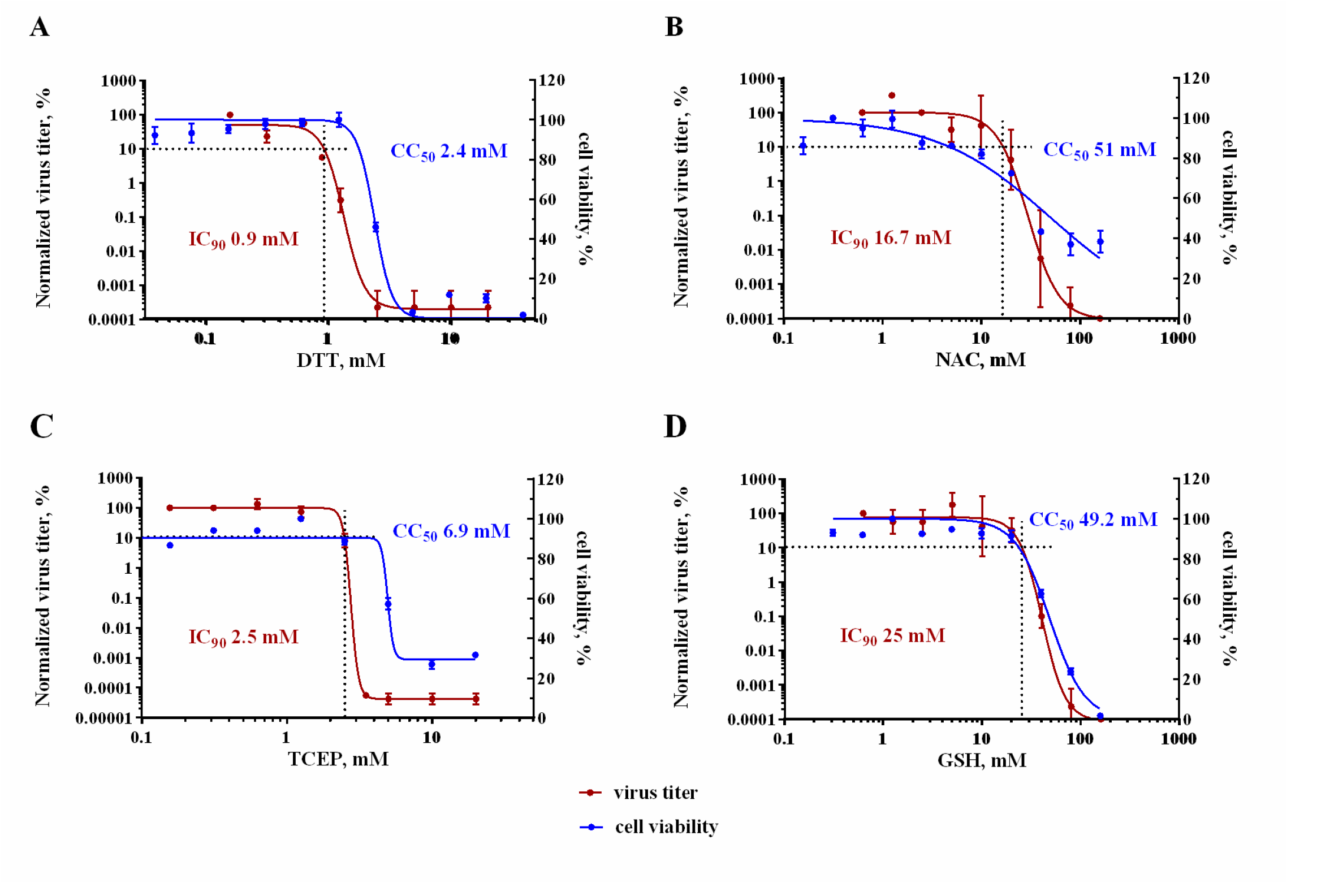
Virus titer and cell viability as a function of the concentration of A) DTT, B) NAC, C) TCEP, and D) GSH. The viral titer, expressed as a log(%) of the normalized TCID_50_/ml, (left *Y*-axis) of SARS-CoV-2 and the viability of Vero’76 cell (%) (right *Y*-axis) were plotted against the dose of the compound (mM) (*X*-axis). The compounds were serially diluted and then added individually to both the cells and the SARS-CoV-2 virus (at an m.o.i. of 0.1) for 1 hour at 37°C. After incubation, the virus-compound mixture was added to the cells and incubated for 1 hour at 37°C. After 1 hour, the mixture was removed, replaced with fresh media containing the compounds and the cells incubated for 48 hours, after which the viral supernatants were harvested and titrated by the TCID_50_ assay. The viability assay was performed in uninfected cells: after 48 hours of incubation with the compounds, the cell viability was assessed using resazurin assay.

We subsequently tested whether DTT, TCEP, NAC and GSH could block viral infection. Viral infectivity assays were conducted by infecting Vero’76 cells with the SARS-CoV-2 and determining the virus titer two days later. Both cells and the virus were preincubated separately for 1 hr with the compounds, then cellular media was discarded and the virus solution was added to the cells for 1 hr. Next, the virus was washed away, while the cells were transferred into fresh media with the compounds for another 48 hr, after which the viral titer in the supernatant was determined.

Out of four compounds tested, DTT and TCEP demonstrated the strongest antiviral effect, with an inhibitory concentration 90% (IC_90_), signifying a 10-fold decrease in the viral titer, of 0.9 and 2.5 mM, respectively (Figure 5A, C). For both compounds, cell viability remained at approximately 100% at IC_90_ concentrations. The decline in cell viability, as evidenced by CC_50_, occurred at compound concentrations exceeding IC_90_ by ∼2.5 times. Thus, both compounds have shown a prominent antiviral effect with the presence of a defined therapeutic window.

For NAC and GSH, the IC_90_ was much higher, 16.7 and 25 mM, respectively (Figure 5B, D), however, their cytotoxicity coincided with the decrease in the viral titer. At both IC_90_concentrations, the cell viability was around 70-80%. The CC_50_ values were 51.0 mM and 49.2 mM for NAC and GSH, respectively. These results suggest that very high and probably unachievable concentrations of these compounds are needed to block viral entry and a very narrow if any therapeutic window measured in the cell culture model.

## Discussion

Here, we provide the first experimental evidence that disulfide-reducing agents can block SARS-CoV-2 infectiousness. Initial molecular modeling of the Spike RBD domain suggested that the reduction of the S–S bond in the surface loop 456-490 may play a paramount role in RBD-ACE2 binding impairment. Addition of reagents capable of S–S reduction altered RBD secondary structure composition and largely decreased its melting temperature bringing it close to physiological range. Additionally, RBD binding to ACE2 was significantly weakened in the presence of DTT and TCEP. Our *in vitro* data on binding impairment correlated well with our cell-based assays, showing the ability of DTT and TCEP to prevent virus penetration and replication in human cells. Our work opens the door to further research on the effect of disulfide reducing agents on the entire architecture of the Spike trimeric ectodomain, conformation destabilization and its membrane fusion activity.

It is not surprising that DTT and TCEP were the most potent in our assays in disrupting the RBD architecture, weakening its interaction with ACE2 and preventing viral infection in comparison to NAC and GSH as the redox potential of monothiols is generally weaker (Chau and Nelson, 1991; Rothwarf and Scheraga, 1992; Gilbert, 1995; Noszál *et al*., 2000; Cline *et al*., 2004). Both the dithiol DTT and the phosphine TCEP are capable of the two-electron reduction of disulfide bonds without the involvement of any other species. DTT and TCEP have similar redox potentials (−330 mV and −290 mV, respectively) and form relatively unreactive oxidized products - a six-membered dithiane ring and a phoshine oxide, respectively (Cleland, 1964; Pullela *et al*., 2006). The mono-thiols NAC and GSH, on the other hand, require the presence of two such thiol-containing molecules to reduce disulfide bonds and would thus be expected to be inherently less effective than DTT and TCEP.

Many toxins and pathogens rely on the proper redox state of their S–S bonds or sulfhydryl groups for host cell entry. For example, the function of the Spike protein of SARS-CoV, which relies on the proper arrangement of disulfide bridges, could be blocked by a DTT concentration somewhere in the 0.5-3 mM range (Lavillette *et al*., 2006). Similarly, the hepatitis C cell entry, dependent on the E2 envelope protein, which also contains multiple S–S bonds, could be blocked by 1 mM DTT (Fenouillet *et al*., 2008). Additionally, the oxidized status of E2 thiol groups is important for evading the production of neutralizing antibodies (Fenouillet *et al*., 2008). Botulinum neurotoxin B is a complex of 2 chains, connected by a single S–S bond. The reduction of this bond by TCEP prior to the neurotoxin internalization prevented the penetration of the active catalytic chain and saved the cells from intoxication (Shi *et al*., 2009). The opposite is also true for other toxins and pathogens – disulfide bond reduction and exposure of sulfhydryl groups promoted the internalization of HIV-1 (Stantchev *et al*., 2012), rotavirus (Calderon *et al*., 2012), Newcastle disease virus (Jain *et al*., 2009) and diphtheria toxin (Ryser HJ *et al*., 1991).

Several opinion articles and reviews have advocated for the plausible role of N-acetyl-cysteine (NAC) in treatment against coronaviral infection (Rangel-Méndez and Moo-Puc, 2020; Poe and Corn, 2020; De Flora *et al*., 2020). Currently, NAC is used as a mucolytic for the reduction of disulfide bonds in mucus (Medici and Radielovic, 1979) and the mitigation of hepatic injury during acetaminophen overdose (Hendrickson, 2019). Various mechanisms have been proposed as to how NAC could help in mitigating SARS-induced conditions, some pointing at the reduction of inflammation and decrease of T cell exhaustion, while others at the improved redox balance, replenishment of reduced glutathione (GSH) and the inhibition of SARS-CoV-2 binding to cells (Silvagno *et al*., 2020; Poe and Corn, 2020; De Flora *et al*., 2020). We would also like to add the low price and high availability of NAC in most countries to this list.

Current usage of NAC in a clinical setting against the coronavirus infection shows controversial results. While in some clinical cases a positive outcome was reported (Ibrahim *et al*., 2020; Alamdari *et al*., 2020), a recent double-blinded clinical study conducted in Brazil did not find a significant difference between placebo and experimental groups (de Alencar *et al*., 2020). More evidence for the effectiveness of NAC in SARS-CoV-2 treatment should be obtained from newly established clinical trials: “Efficacy of N-Acetylcysteine (NAC) in Preventing COVID-19 from Progressing to Severe Disease,” identifier NCT04419025 and “A Study of N-acetylcysteine in Patients with COVID-19 Infection,” NCT04374461 (clinicaltrials.gov).

Nevertheless, our data show that NAC is likely to be ineffective against SARS-CoV-2 as a cell entry inhibitor. Pharmacological studies of NAC have shown that during a standard regimen with a total dose of 300 mg/kg, administered over 20 hours with the initial loading of 150 mg/kg over the first 15 min, the plasma concentration of NAC achieves on average 554 mg/L (3.4 mM), range: 304–875 mg/L, after initial loading, but then rapidly falls to 35 mg/L (0.21 mM). The half-life of NAC was around 6 hours (Prescott *et al*., 1989). According to our results, even the peak concentration is not sufficient to prevent viral infection as we did not observe a significant decrease in the viral titer in NAC concentrations as high as 10 mM. Nevertheless, NAC may still be a plausible medication against SARS-CoV-2 infection as it may exert its action through other diverse mechanisms postulated in plenty by the scientific community (Silvagno *et al*., 2020; Poe and Corn, 2020; De Flora *et al*., 2020).

A high dose 48-hour regimen was developed for NAC in which the initial loading of 140 mg/kg is followed by 70 mg/kg every 4 hours for 12 doses in total with the cumulative dose of 980 mg/kg (Heard *et al*., 2014). NAC can also be administered through nebulization, in which 4 ml of 20% solution is given 3-4 times/day (Gallon, 1996), which should produce high concentrations of NAC in the lungs. Whether any of these regimens will be effective against SARS-CoV-2, or whether even higher dose regimens have to be developed, is not yet known. Another variable is the proper duration of NAC treatment. One successful case of bronchoscopy lavage with 10-15 ml of NAC inhalation solution for an elderly patient was reported (Liu *et al*., 2020).

DTT has been tested in humans as a medication against amyotrophic lateral sclerosis at a 1-2 gr daily dose, which is still sub-mM range. A broad range of side effects was observed, although no major life-threatening conditions were detected (Vyth *et al*., 1996). Another attempt to use DTT against cystinosis in pediatric patients did not detect major side effects during 31 patient-treatment months with a dose of 75 mg/kg (Depape-Brigger *et al*., 1977). The mouse lethal dose 50% (LD_50_) of DTT was higher for the D-versus L-enantiomer, 255 mg/kg versus 179 mg/kg, with only the D-enantiomer having protective properties against X-ray radiation (Carmack *et al*., 1972). These LD_50_ values are of a similar low mM range to the CC_50_ value that we have observed in cell-based assays.

GSH is a natural metabolite present in the cytosol at a concentration range of 0.5 to 10 mM (Kosower and Kosower, 1978) and in plasma at 1000 times lower concentrations (Jones *et al*., 2000). Thus, anti-SARS therapy could aim at increasing plasma GSH concentration. A recent case report indicated that administration of 2 g of GSH improved SARS-CoV-2-induced dyspnea within one hour of administration for two patients in New York (Horowitz *et al*., 2020). Our results show that GSH is not very potent against SARS-CoV-2 and that its antiviral effect is likely due to cytotoxicity to the host cells after 48 hours of incubation. However, the short-term administration of this compound might result in a decrease in viral titer without significant change in cellular toxicity.

While TCEP is not currently considered as a potential medication, its toxicity seems to be low with an oral LD_50_ for rat of 3500 mg/kg, according to the MSDS. 100 µM TCEP was shown to help retinal ganglion cells in their long-term survival after axotomy (Geiger *et al*., 2002). TCEP was benign to glial cells for up to 3 mM concentration (Cha *et al*., 2018). Similarly, 1-2 mM TCEP concentration had an insignificant impact on the viability of SHSY-5Y neuroblastoma cell line (Shi *et al*., 2009). Our data showed that TCEP started to develop toxicity to the cells at concentrations >5 mM, although cells were not completely dead even at 20 mM concentration.

Of the four compounds we tested, DTT and TCEP, both possessing a strong redox potential, showed significant antiviral activity in the high µM - low mM range without causing marked cytotoxicity. Thus, DTT and TCEP serve as proof of principle that disulfide-reducing agents could be used to stop SARS-CoV-2 infection as cell entry inhibitors.

Overall, disulfide-reducing agents should be studied further for their potential against SARS-CoV-2 and their effect on human health for the risk of disrupting biologically relevant S–S bridges in antibodies and other human proteins. These studies should be accompanied by pharmacological, toxicological and dosing regimen elucidation. In the meantime, more disulfide-reducing agents can be developed to help the world cope with sudden and unexpected coronavirus outbreaks. The alternatives to N-acetyl-cysteine include approved medications S-carboxymethyl-L-cysteine (Mitchell and Steventon, 2012) and erdosteine (Cazzola *et al*., 2020) among others, along with an experimental compound I-152, which is a prodrug with conjugated NAC and β-mercaptoethylamine (Crinelli *et al*., 2019). A recent manuscript submitted to BioRxiv explores the impact of 8 approved thiol-containing drugs on SARS-CoV-2 infectivity and cell entry with the conclusion that cysteamine and WR-1068 are the most potent inhibitors, exhibiting their action in high µM – low mM range (Khanna *et al*., 2020). Many other dithiol-based compounds, such as dithioerythritol (DTE) (Vyth *et al*., 1996) and dithiobutylamine (DTBA) (WO 2013/123382 Al), and phosphine-based compounds, such as monomethyl-tris(2-carboxyethyl)phosphine (mmTCEP) (Cline *et al*., 2004) or tris(3-hydroxypropyl)phosphine (THPP) (McNulty *et al*., 2015), are available for further investigation.

Because the evolution of coronaviruses proceeds in the wild, new coronaviruses emerging in the future will likely rely on the S–S bonds for the Spike protein architecture. Thus, disulfide-reducing agents may become the first line of defense against any coronavirus outbreak. If an appropriate concentration cannot be achieved systemically, it may still be possible to achieve it locally in the lungs through nebulization. In this case, disulfide reducing agents may serve not only as a medication to clear patient lungs of a coronavirus but also as a preventive measure for risk groups, inhibiting inhaled virus particles from entering the cells. The sensitivity of the new coronavirus to NAC and any other disulfide-reducing agent approved as a medication at that time could be tested within weeks in the laboratory environment, followed by an accelerated limited clinical trial.

## Acknowledgments

- We gratefully acknowledge the use of instruments at the Protein Characterization and Crystallization Facility, College of Medicine, the University of Saskatchewan funded by the Canada Foundation for Innovation. We thank the members of Information and Communications Technology, University of Saskatchewan and specifically Dr. Olivier Fisette for access to the Research Computing Cluster resources and help with Gromacs molecular dynamics software. We thank Dr. Oleg Dmitriev, Department of Biochemistry, Microbiology and Immunology, College of Medicine, the University of Saskatchewan and members of his laboratory for support and help with equipment.
- This work was supported by the Canadian Institutes of Health Research grant GSP-48370 (M.C.), The Natural Sciences and Engineering Research Council RGPIN-2016-05810 (I.J.P.) and RGPIN-2019-05351 (G.N.G.), the University of Saskatchewan, and the Canada Research Chairs Programs (I.J.P., G.N.G., M.C.).

