## Supplemental Figures 1-3 for "Spike protein disulfide disruption as a potential treatment for SARS-CoV-2"

THIS PDF INCLUDES:

Supplemental Figures 1-3

Figure S1. The final stage of purification of A) RBD-tev-his$_6$ and B) ACE2. The figure shows a gel-filtration curve with Superdex 75 16/600 column followed by the SDS-PAGE image of the consecutive fractions of the peak (marked by an arrow). RBD has a smeared appearance on the gel due to glycosylation.

A

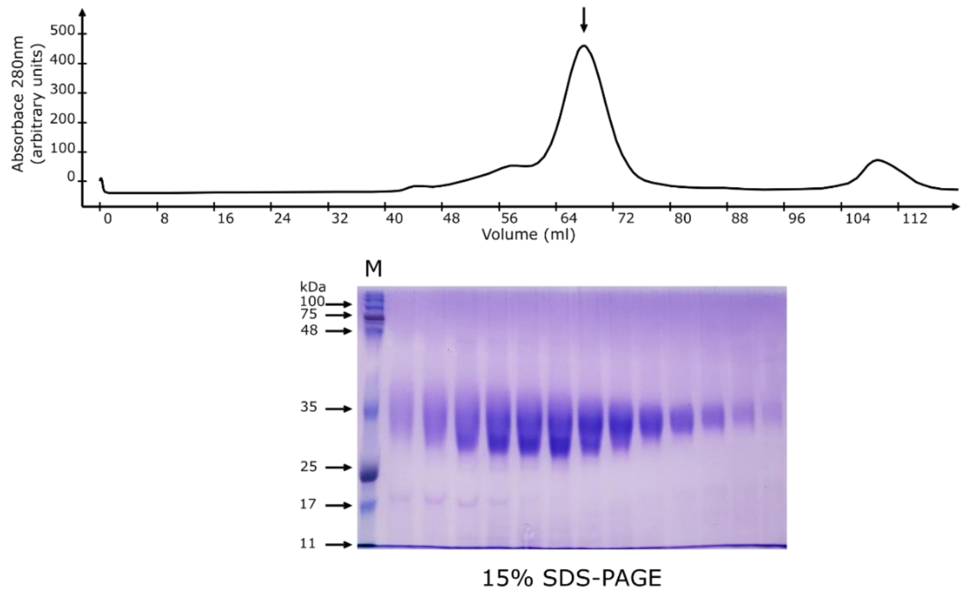


B

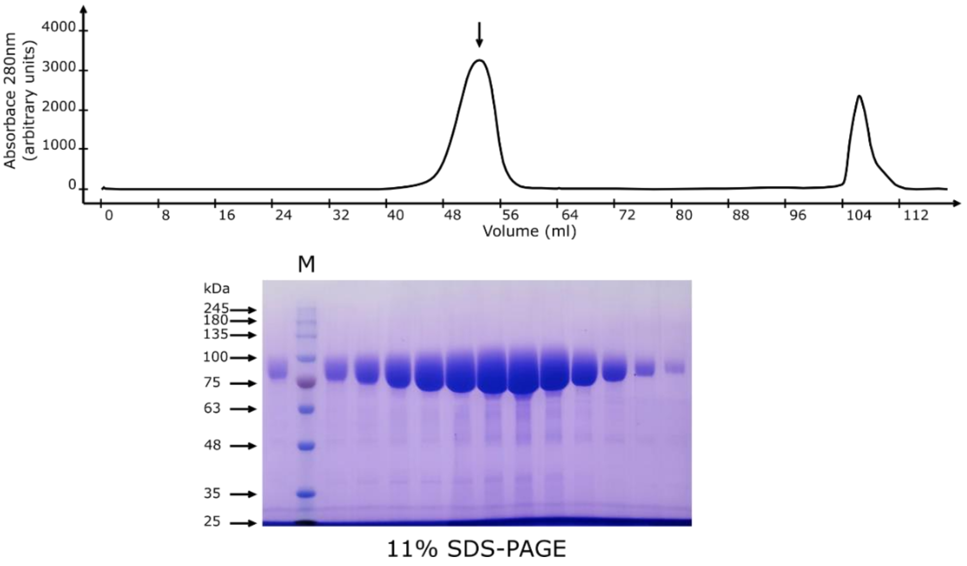

Figure S2. Comparison of circular dichroism spectra of RBD. RBD was preincubated for 1 hour at 37°C in the presence of disulfide-reducing agents at 2.5 mM concentration and cooled down to RT for 30 min. Spectra were acquired at RT. The spectrum of RBD, never exposed to elevated temperatures and acquired at room temperature (RT), is provided for reference.

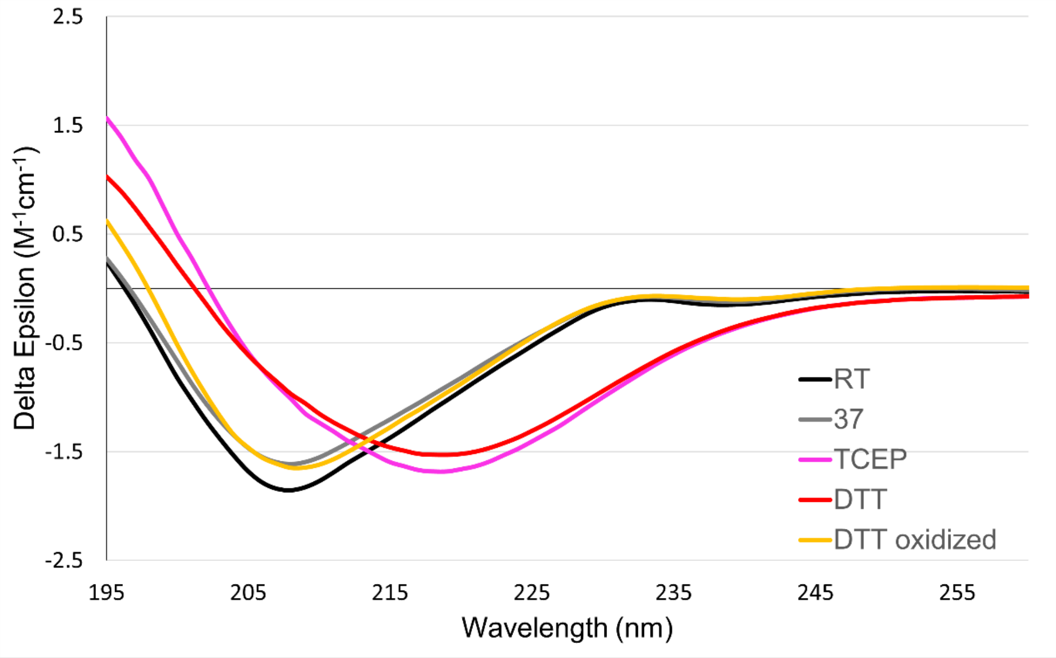

Figure S3. Brigthfield microscopy (20x) pictures of Vero’76 cells in the presence of (A) DTT, (B) NAC, (C) TCEP and (D) GSH at concentrations close to their corresponding $CC_{50}$ values.

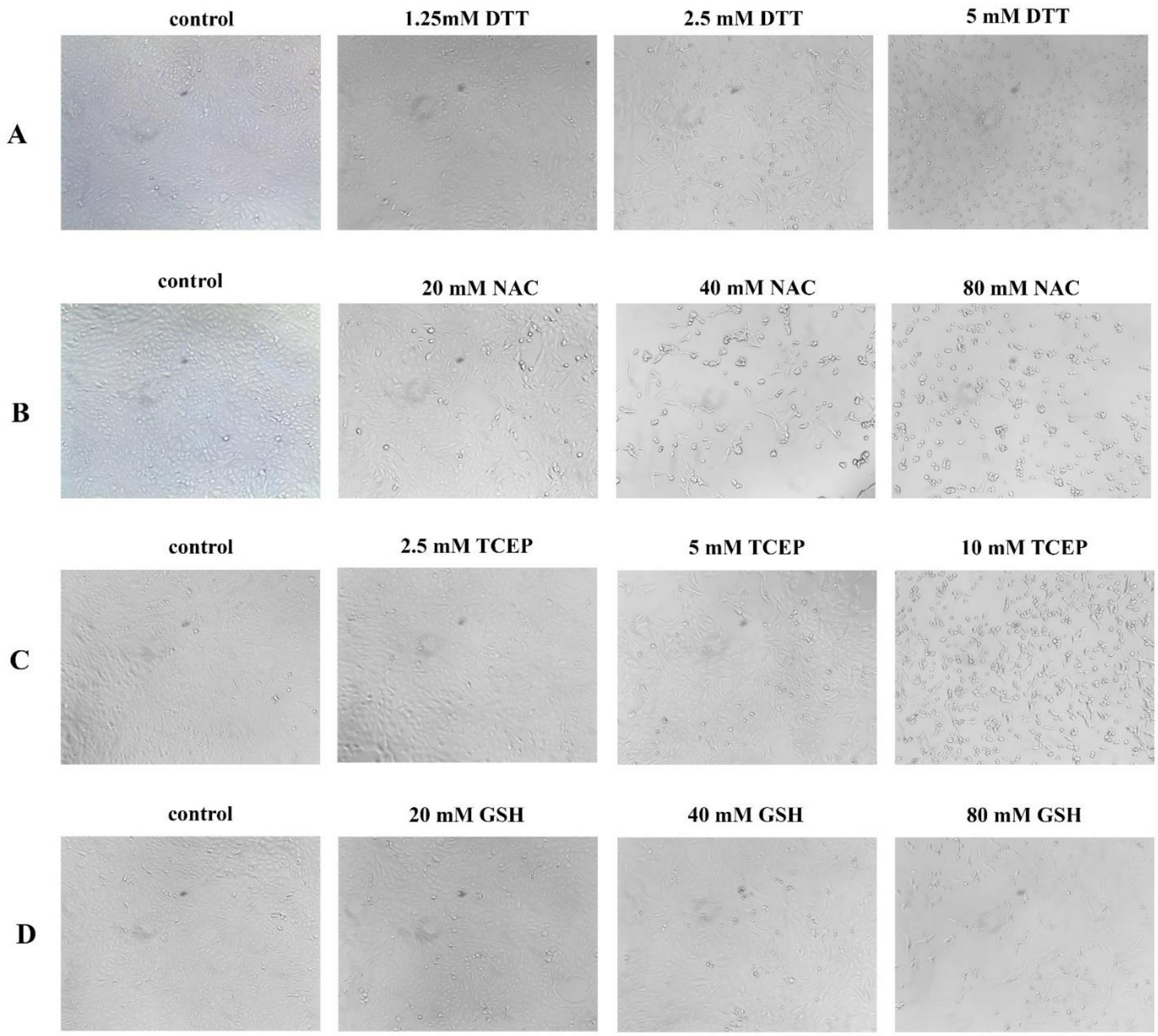
